# Gene expression variability across cells and species shapes innate immunity

**DOI:** 10.1101/137992

**Authors:** Tzachi Hagai, Xi Chen, Ricardo J Miragaia, Tomás Gomes, Raghd Rostom, Natalia Kunowska, Valentina Proserpio, Giacomo Donati, Lara Bossini-Castillo, Guy Naamati, Guy Emerton, Gosia Trynka, Ivanela Kondova, Mike Denis, Sarah A Teichmann

**Affiliations:** Wellcome Trust Sanger Institute, Wellcome Genome Campus, Hinxton, Cambridge, CB10 1SD, UK; EMBL- European Bioinformatics Institute, Wellcome Genome Campus, Hinxton, Cambridge, CB10 1SD, UK; Centre of Biological Engineering, University of Minho, Braga, Portugal; Department of Life Sciences and Systems Biology, University of Turin, Via Accademia Albertina 13, 10123 Torino, Italy; Division of Pathology and Microbiology, Biomedical Primate Research Centre, 2280 GH Rijswijk, the Netherlands; Research Department. Public Health England, National Infection Service, Porton Down, Wiltshire SP4 0JG, UK; Theory of Condensed Matter Group, Cavendish Laboratory, University of Cambridge, JJ Thomson Avenue, Cambridge CB3 0HE, UK

## Abstract

As the first line of defence against pathogens, cells mount an innate immune response, which is highly variable from cell to cell. The response must be potent yet carefully controlled to avoid self-damage. How these constraints have shaped the evolution of innate immunity remains poorly understood. Here, we characterise this programme’s transcriptional divergence between species and expression variability across cells. Using bulk and single-cell transcriptomics in primate and rodent fibroblasts challenged with an immune stimulus, we reveal a striking architecture of the innate immune response. Rapidly diverging genes, including cytokines and chemokines, also vary across cells and have distinct promoter structures. Conversely, genes involved in response regulation, such as transcription factors and kinases, are conserved between species and display low cell-to-cell variability. We suggest that this unique expression pattern, observed across species and conditions, has evolved as a mechanism for fine-tuned regulation, to achieve an effective but balanced response.

## Introduction

The innate immune response is a cell-intrinsic defence program that is rapidly upregulated upon infection. It acts to inhibit pathogen replication while signalling the pathogen’s presence to other cells. This programme involves modulation of several cellular pathways, including production of antiviral and inflammatory cytokines, changes in translation and metabolism, upregulation of numerous genes functioning in cell defence and pathogen restriction, and the induction of regulated cell death^1,2^.

An important characteristic of the innate immune response is the rapid evolution that many of its genes have undergone along the vertebrate lineage. Comparative analyses across genomes have highlighted immune genes as having extremely rapid coding sequence divergence^3,4^. This is often attributed to pathogen-driven selective pressure^5-8^.

Another hallmark of the innate immune response is its high level of heterogeneity among responding cells: various studies have shown that cells display extensive cell-to-cell variability in response to pathogen infection^9,10^ and exposure to pathogen-associated molecular patterns (PAMPs)^11,12^. The functional importance of this cell-to-cell variability in response is unclear.

These two characteristics – rapid divergence in the course of evolution and high cell-to-cell variability – seem to be at odds with the strong regulatory constraint imposed on the innate immune response: the need to execute a well-coordinated and carefully balanced programme, to avoid long-term tissue damage and pathological immune conditions^13-16^. How this tight regulation is maintained despite rapid evolutionary divergence and high cell-to-cell variability remains an open question, central to our understanding of the innate immune response and its evolution.

Here, we combine three genomics approaches to comprehensively characterize the transcriptional changes between species and among individual cells in their innate immune response (Fig 1A). We use population (bulk) transcriptomics to investigate transcriptional divergence between species, and single cell transcriptomics to estimate cell-to-cell variability in gene expression. Through integration of these two approaches with chromatin immunoprecipitation followed by sequencing (ChIP-seq) and sequence analyses, we study how changes in the expression of each gene between species and across cells relate to the evolution of its promoter and coding sequence. Finally, we examine the relationship between divergence in sequence and expression and constraints imposed by host-pathogen interactions, to elucidate the evolutionary architecture of the innate immune response.

**Figure 1:**
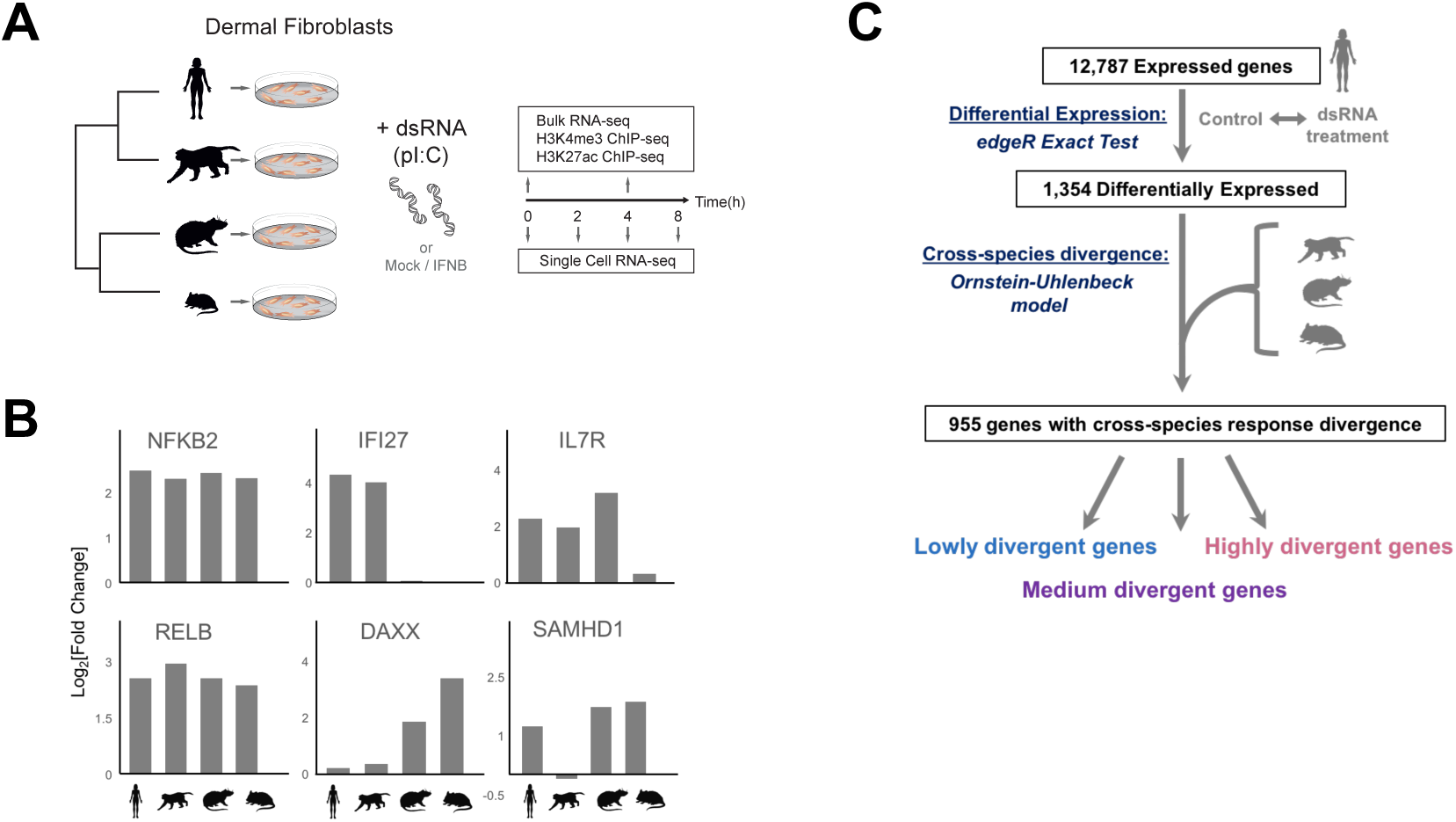
Transcriptional divergence in the dsRNA response. **(A)** Study design: Primary dermal fibroblasts from four species - stimulated with dsRNA, or controls (unstimulated cells, mock transfection), or an alternative stimulation (IFNB). Samples were collected for population (bulk) and single-cell RNA-seq and for ChIP-seq. **(B)** Fold change in response to dsRNA in six example genes across the four species: NFKB2, RELB, IFI27, DAXX, IL7RA and SAMHD1. **(C)** Estimating each gene’s level of cross-species divergence in transcriptional response: (1) Using edgeR, the fold change in expression in response to dsRNA (in comparison to mock transfection) was assessed for each gene in each species. (2) Differentially expressed genes were identified using the edgeR exact test (FDR-corrected q-value<0.01), out of which 955 have one-to-one orthologs across the four studied species. (3) Ornstein-Uhlenbeck models were fitted to the fold change data for each gene, using the expression changes of the four orthologs and the known phylogeny (see Methods). Genes were grouped into ‘low’, ‘medium’ and ‘high’ divergence for subsequent analysis.

Our experimental setup uses primary dermal fibroblasts, which are commonly employed in immunological and virological studies, and provide an excellent *in vitro* system to study the evolution of the innate immune response across species. We compare the response of dermal fibroblasts from primates (human and rhesus macaque) and rodents (mouse and rat) to poly I:C, a synthetic dsRNA. Poly I:C, a primary PAMP, is frequently used to mimic viral infection as it rapidly elicits an innate immune response^17^.

Using this model, we decipher both transcriptional divergence and cell-to-cell variability, allowing us to probe the evolutionary mechanisms underlying the heterogeneity in the innate immune response.

## Results

### Estimating transcriptional divergence in the dsRNA-response across species

To compare the transcriptional response across the four species (human, macaque, rat and mouse), we generated bulk RNA-sequencing data from the fibroblasts of each species after stimulation with dsRNA (poly I:C) for 4 hours, along with respective controls (see Fig 1A, Table S1 and Methods for detailed experimental design and controls used).

In all species, we observe rapid upregulation of expected antiviral and inflammatory genes, including IFNB, TNF, IL1A and CCL5, following dsRNA treatment (see also Table S2 for upregulated pathways). Correlation analysis between species shows a similar transcriptional response (Fig S1A-C), as reported in other immune contexts^18-20^. Furthermore, the response tends to be more conserved between closely related species (Fig S1I), similarly to other expression programmes^21-25^

We characterized the differences between species in response to dsRNA for each gene, using this cross-species bulk transcriptomics data. While some genes, such as the NF-*κ*B subunits RELB and NFKB2, have a similar response across species (Fig 1B), other genes respond differently between the primate and the rodent clades. For example, IFI27 (a restriction factor against numerous viruses) is strongly upregulated in primates but not in rodents, while DAXX (an important antiviral transcriptional repressor) exhibits the opposite behaviour (Fig 1B). Conversely, some genes have species-specific response patterns. For example, IL7RA (a cytokine receptor) is upregulated in response to dsRNA in human, macaque and rat, but not in mouse, in agreement with previous observations from LPS stimulation of human and mouse macrophages^19^ (see additional examples in Figs 1B and S2).

To quantify divergence in transcriptional response to dsRNA between species, we focused on genes that are differentially expressed in human in response to dsRNA (see Methods). For simplicity, we will refer to these genes as “dsRNA-response genes” (Fig 1C). There are 955 such genes with one-to-one orthologs across the species.

For each gene, we fitted models that incorporate selection and genetic drift to the changes in gene expression after stimulation across the four species (Ornstein–Uhlenbeck model^26^ - hereafter OU model, see Fig 1C and Methods for additional details). This procedure provides a measure of divergence in transcriptional response to dsRNA between species that is comparable across all genes (hereafter “response divergence”).

For subsequent analyses, we split the 955 dsRNA-response genes, based on their level of response divergence, into three groups: (1) highly divergent dsRNA-response genes (the top 25% genes with the highest divergence values in response to dsRNA across the four studied species), (2) lowly divergent dsRNA-response genes (the bottom 25%), and (3) genes with medium divergence across species (the middle 50%) (Fig 1C).

We confirmed the results of the OU divergence model by observing agreement between these values and divergence measures obtained by a different model (linear models between species - Fig S3). We then performed ChIP-seq of stimulated and unstimulated cells in all species to profile two histone marks: H3K4me3 and H3K27ac. Both of these marks are present at active promoters, while H3K27ac without H3K4me3 is associated with active enhancers. Our analysis showed that divergence in gene expression corresponds to divergence of active promoters and enhancers across the four species (Fig S4).

### Rapidly diverging genes have a distinct promoter architecture

Next, we tested whether divergence in transcriptional response is reflected in promoter sequence conservation and architecture. For this, we compared the conservation of the 500 bases upstream of the transcription start site (TSS) in highly divergent dsRNA-response genes with conservation of the corresponding region in medium and lowly divergent genes (Fig 2A).

Interestingly, genes that highly diverge in response show higher sequence conservation in this region. Thus, despite having a relatively conserved promoter sequence, these genes dramatically differ in their expression levels in response to dsRNA between species. This surprising discordance may be related to the fact that promoters of highly and lowly divergent genes have different architectures, which impose different constraints on promoter sequence evolution^19,27,28^. Promoters containing TATA-box elements tend to have most of their regulatory elements in regions immediately upstream of the TSS. These promoters are thus expected to be more conserved. The opposite is true for CpG island (CGI) promoters^27^. Indeed, we found that TATA-box elements tend to occur in promoters of highly divergent genes, while CGIs occur in more conserved genes (Fig 2B). Thus, a promoter architecture enriched with TATA-boxes and depleted of CGIs is associated with higher transcriptional divergence, while entailing higher sequence conservation upstream of these genes. Conversely, genes with CGI promoters tend to be transcriptionally conserved^19,27,28^.

**Figure 2:**
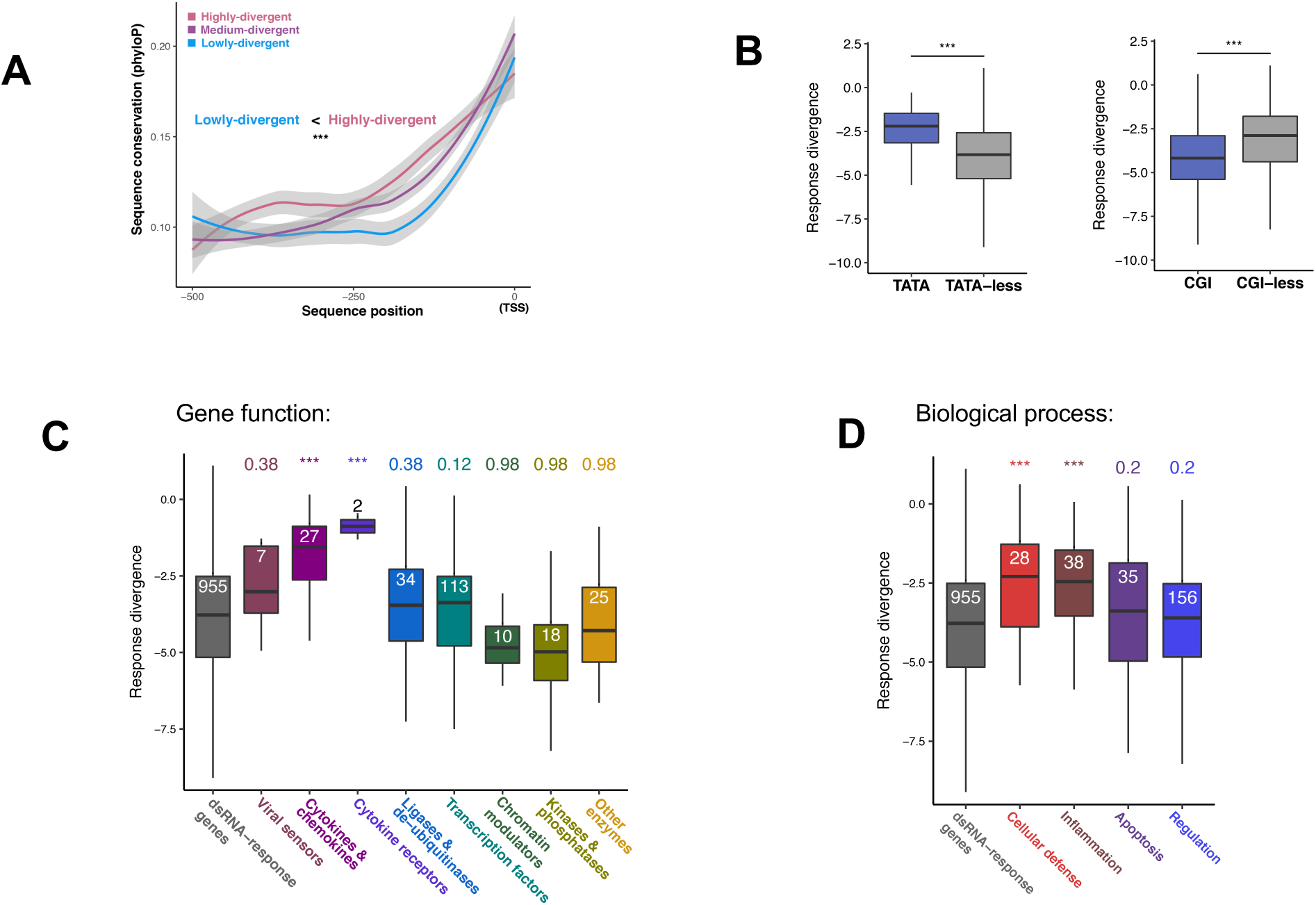
Transcriptional response divergence, promoter architecture and gene function. **(A)** Promoter sequence conservation and transcriptional response divergence: Sequence conservation values are estimated with phyloP7 (higher values indicate higher conservation) for 500bp upstream the transcription start site (TSS). Mean conservation values each bp in this region upstream of the TSS are shown for highly-, medium- and lowly-divergent genes. Genes that are highly-divergent have higher sequence conservation (p=3.8x10-7, two-sample Kolmogorov-Smirnov test). **(B)** Comparison of divergence in response of genes with and without a TATA-box (left) and CpG Islands (CGIs) (right). Divergence in response values (y-axis) are taken from the Ornstein-Uhlenbeck model. Empirical p-values are shown. **(C-D)** Distributions of divergence values of all 955 dsRNA-response genes and different functional subsets of this group (see Methods for details on each of the functional subsets): (C) Division by gene’s molecular function; (D) Division by biological process. FDR-corrected empirical p-values are shown above each group; group size is shown inside each box. Values of response divergence are estimated as 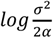 where α and σ are the coefficients for selection and drift (respectively) from a fitted Ornstein-Uhlenbeck model (see Methods). Higher values (in y-axis in B-D) indicate faster response divergence. (For all panels: *p<0.05, **p<0.01,***p<0.001).

### Cytokines and chemokines diverge more rapidly than other responding genes

We next asked whether dsRNA-response genes from different functional classes diverge in their transcriptional response at different evolutionary rates. For this, we divided the 955 dsRNA-response genes into different categories, based on their function (such as cytokines, transcriptional factors, kinases) or based on the processes in which they are known to be involved (such as apoptosis or inflammation). We observed that genes that are related to cellular defence and inflammation – most notably cytokines, chemokines and their receptors – tend to diverge in response significantly faster than genes involved in apoptosis or immune regulation (chromatin modulators, transcription factors, kinases and ligases) (Fig 2C-D). Similar findings regarding divergence rates of different functions of responding genes were observed in response to direct IFNB treatment, which we used as an additional immune challenge for comparison (Fig S5).

In summary, we have identified a group of genes that have undergone relatively fast divergence in their transcriptional response to dsRNA between the four studied species. These genes have a distinct promoter architecture. Furthermore, these genes have specific functions in innate immunity related to cellular defence and inflammation, including cytokines, chemokines and their receptors. In contrast, dsRNA-induced genes involved immune response regulation (such as kinases and transcription factors) are more conserved in their transcriptional response.

### Rapidly diverging genes have high cell-to-cell variability in expression

Previous studies have shown that the innate immune response displays high variability across responding cells^29,30^. However, the relationship between cell-to-cell transcriptional variability and response divergence between species is not well-understood.

To study gene expression heterogeneity across cells, we used single-cell RNA-seq to profile the transcriptomes of hundreds of individual cells from all species in a time course of 0,2,4 and 8 hours after dsRNA stimulation. We estimated cell-to-cell variability quantitatively using an established measure for variability: “distance to median” (DM, see Methods)^31^. This measure has been previously used to infer variability from single-cell RNA-seq data^32,33^, and accounts for the confounding factor of gene expression level (Fig S6).

We determined the relationship between divergence across species and cell-to cell variability for the 955 dsRNA-response genes (Fig 3A). Strikingly, we found a clear trend where genes that are highly divergent in response between species are also more variable in expression across individual cells within a species. The relationship between rapid divergence and high cell-to-cell variability holds in each species and each stimulation time point (Fig S7).

We confirmed that these results are not biased by particular computational or experimental settings (Figs S8-9): (1) by repeating the analyses with a second computational approach - BASiCS (a novel approach to infer variability that discerns biological variability from technical noise by utilizing observed levels of synthetic spiked-in RNA molecules^34^), (2) by repeating the analysis with a second biological replicate from each species, and (3) by profiling thousands of human cells that were processed using a different experimental technique (Fig S8). Furthermore, we confirmed that cell-to-cell variability in response does not stem from different overall levels of response (Fig S10).

Thus, the set of genes rapidly diverging in response to dsRNA across species also displays high cell-to-cell variability in expression within a species - a relationship that is consistent across all species and stimulation time points.

**Figure 3:**
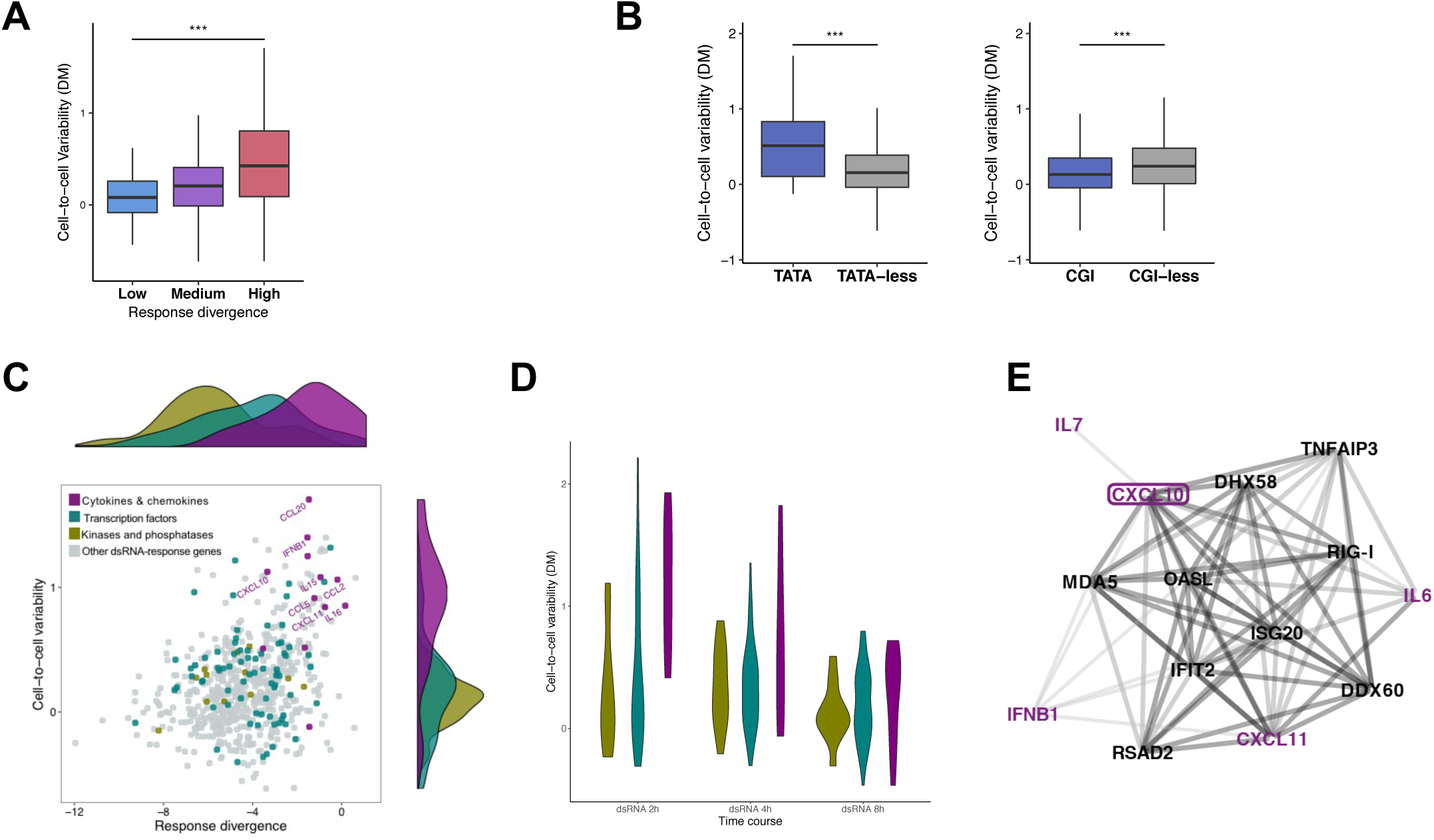
Cell-to-cell variability in the dsRNA response. **(A)** A comparison of divergence in response (measured in bulk transcriptomics across four species) and transcriptional variability in gene expression between individual cells (measured in single-cell transcriptomics in human cells) in 955 dsRNA-response genes. In both measures, cells were stimulated with dsRNA for 4 hour. Cell-to-cell variability was estimated using DM (higher values indicate higher levels of variability). Genes are split into highly-, medium- and lowly-diverging genes based on their level of response divergence (x-axis, as described in Figure 1C). Cell-to-cell variability values of highly-divergent genes were compared with lowly-divergent genes using Mann-Whitney test (*p<0.05, **p<0.01, ***p<0.001). **(B)** Comparison of cell-to-cell variability of genes with and without a TATA-box (left) and CpG Islands (CGIs) (right); empirical p-values are shown. Cell-to-cell variability values (y-axis) are taken from the DM estimations of human cells stimulated with dsRNA for 4 hours. **(C)** A scatter plot showing divergence in response to dsRNA across species (x-axis) and transcriptional cell-to-cell variability in human cells following 4 hours of dsRNA treatment (y-axis), of 955 dsRNA-response genes. Genes from three functional groups - cytokines, transcription factors and kinases - are coloured, and some cytokines’ names are shown. The distributions of divergence values of each of the three functional groups are shown above the scatter plot (with colours matching the scatter plot). The distributions of cell-to-cell variability values of these groups are shown to the right. **(D)** Violin plots showing the distribution of cell-to-cell variability values of cytokines, transcription factors and kinases following dsRNA treatment for 2,4 and 8 hours (gene groups are colours as in C). **(E)** A network showing 14 genes that correlate in expression across cells with the chemokine CXCL10, in at least two species, following dsRNA treatment. Here we include genes correlating with CXCL10 that are either cytokines (colored in purple) or genes that are known to be associated with cytokine upregulation (genes with GO terms: “positive regulation of innate immune response” or “type I interferon signalling pathway”). See Figure S14 for the network that includes all genes that were found to correlate in expression with CXCL10 in at least two species (colours of edges, from light to dark grey, reflect the number of species in which this pair of genes was found to be correlated).

### High cell-to-cell variability in expression is associated with unique promoter architecture

Next, we examined the relationship between the presence of promoter elements – CpG islands (CGIs) and TATA-boxes - and a gene’s cell-to-cell variability. We observed that genes that are predicted to have a TATA-box in their promoter have significantly higher transcriptional variability, while CGI-genes tend to have significantly lower variability (Fig 3B). Thus, both transcriptional variability between cells (Fig 3B) and transcriptional divergence between species (Fig 2B) are associated with the presence of specific promoter elements. TATA-box presence is associated with high divergence and variability, while CGIs occur in promoters of genes with conserved transcriptional response and low cell-to-cell variability.

### Cytokines and chemokines display high levels of cell-to-cell variability across species and conditions

We subsequently investigated the transcriptional cell-to-cell variability of three groups of dsRNA-response genes with different functions: cytokines and chemokines (hereafter “cytokines”), transcription factors, and kinases and phosphatases (hereafter “kinases”). Cytokines rapidly diverge in transcriptional response, while the genes in the other two groups are relatively conserved (Fig 2C). We compared response divergence across species with levels of cell-to-cell variability for each dsRNA-response gene (Fig 3C). In contrast to kinases and transcription factors, many cytokines display relatively high levels of cell-to-cell variability. This high variability is a manifestation of cytokine expression in a small subset of cells (Fig S11). The expression of cytokines in only a subset of cells within a responding cell population has been reported for several cytokines in various innate and adaptive immune scenarios^30^. For example, IFNB is expressed in only a small fraction of cells infected with different viruses or challenged with various stimuli^9,12,35^. Here, we observed that cytokines from several different families (e.g. IFNB, CXCL10, CCL2) are upregulated in a small fraction of the stimulated cells, and thus exhibit high levels of expression heterogeneity between cells.

When looking at the progression of the response to dsRNA along a time course of 2, 4 and 8hrs, we observe that the cell-to-cell variability of cytokines is relatively high at all time points, in comparison with kinases and transcription factors that display relatively low variability (Fig 3D). This pattern is similar across all species (Fig S12). Thus, the high variability of cytokines and their expression in a small fraction of stimulated cells across all time points is evolutionarily conserved (Figs S11-12).

### Co-expression of cytokines with upstream regulators is conserved across species

When analysing cytokine expression across dsRNA-stimulated human cells, we found a group of 13 cytokines and chemokines that tend to be co-expressed in the same cells (Fig S13A). This set includes some of the most heterogeneously expressed cytokines, such as IFNB1, CXCL10, CXCL11, CCL2 and CCL5 (Fig S13B). This raises the possibility that the expression of these cytokines is coordinated.

To further investigate this, we calculated the correlations between expression of these 13 cytokines and other expressed genes across dsRNA-stimulated cells. We observed that the set of genes correlating with these 13 cytokines (genes with >0.3 R-Spearman correlation) is enriched in GO terms related to the innate immune response, such as “Defence response to virus” and “Type I interferon signalling pathway” (Table S3A). Next, we repeated the same correlation analysis in macaque, mouse and rat cells. We showed that the set of orthologs of genes identified in human as correlating with cytokines is significantly co-expressed with cytokines in the other species (see Methods). Among the genes co-expressed with the 13 cytokines in human, 25 genes show conservation of a high correlation with the same cytokines across all four species (Table S3B). This set is enriched with genes known to be involved in cytokine regulation.

As an example, we identified the genes whose expression is correlated with the chemokine CXCL10 in at least two species (Figs 3E and S14). This set of genes includes 4 cytokines that are co-expressed with CXCL10, and 10 genes that are known regulators of the innate immune response and cytokine production. Examples of these genes include the viral sensors and dsRNA binding proteins MDA5 and RIG-I, and their regulator and binding partner DHX58. This is in agreement with previous findings that suggest that IFNB expression is limited to cells where important upstream regulators are expressed at high enough levels^9,12,35^. Here, we showed that this phenomenon of co-expression with upstream regulators applies to a wider set of cytokines and is conserved across species.

In summary, we observe that cytokines are expressed in a small subset of cells, giving rise to high cell-to-cell variability in transcriptional response to dsRNA. This pattern is seen across species and different stimulation time points, and is associated with expression of genes known to be upstream regulators of cytokine production.

### Transcriptionally divergent genes also evolve rapidly in protein sequence

Above, we characterised genes that rapidly diverge in response to dsRNA, and showed that these genes are also variable in expression across cells. Many immune genes, including some cytokines and their receptors, are known to evolve rapidly in coding sequence compared to all other genes^3,6^. However, whether divergence in transcriptional response in immune genes is related to divergence in coding sequence is not known.

This prompted us to examine the relationship between divergence in response and divergence in coding sequence. For these analyses, we used the set of 955 dsRNA-response genes, and assessed the coding sequence evolution in the three subsets of lowly, medium and highly divergent genes (as defined in Fig 1C).

We compared the rate at which genes evolved in their coding sequences with their response divergence by considering the ratio between non-synonymous (dN) to synonymous substitutions (dS). We observed that genes that rapidly evolve in transcriptional response have higher coding sequence divergence (higher dN/dS values) than dsRNA-response genes with low response divergence (Fig 4A).

**Figure 4:**
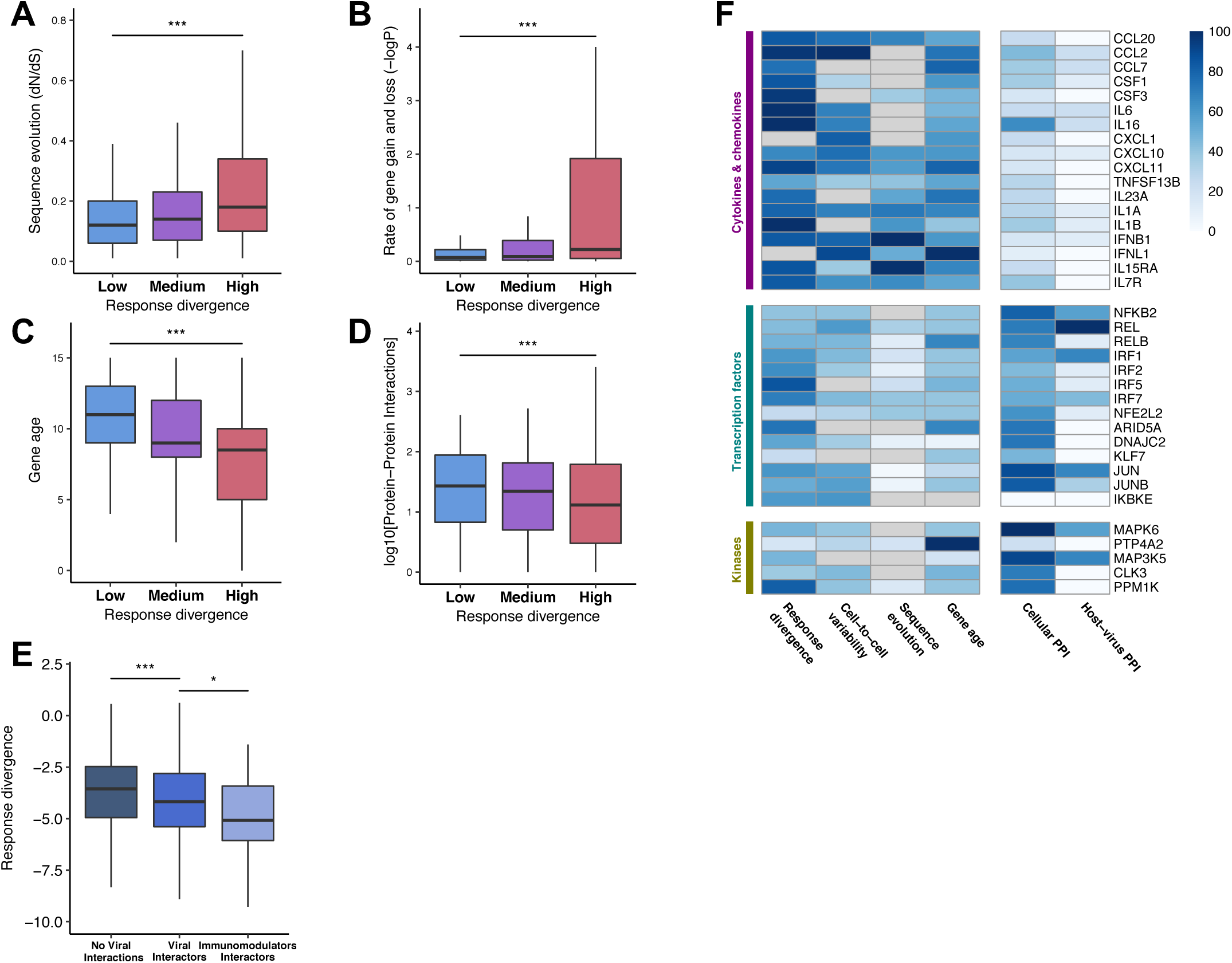
Correspondence of response divergence and other evolutionary modes. Comparing transcriptional divergence in response to dsRNA with other evolutionary mechanisms: In A-D 955 dsRNA-response genes are divided by level of response divergence into three groups: highly divergent genes (top 25% most diverging genes, in pink), lowly divergent genes (25% least diverging genes, in blue), and medium divergent genes (in purple) (as in Fig 1C). In each panel a different measure is shown in the y-axis for each of the three groups (e.g. – coding sequence divergence in panel A). A a statistical comparison (Mann-Whitney test) is made between the distribution of the tested measure’s values in highly and lowly divergent genes. (*p<0.05, **p<0.01, ***p<0.001.) **(A)** Coding sequence divergence, as measured using dN/dS values across a clade of 29 mammals. Higher dN/dS values denote faster coding sequence evolution. **(B)** Rate at which genes are gained and lost within the gene family across the vertebrate clade (plotted as –logP). Higher values denote faster gene gain and loss rate. **(C)** Evolutionary age (estimated with Panther7 phylogeny and Wagner reconstruction algorithm). Values denote the branch number with respect to human (distance from human in the phylogenetic tree), higher values indicate older age. **(D)** Number of known physical interactions with other cellular proteins. **(E)** Distribution of transcriptional response divergence values among dsRNA-response genes that do not interact with viruses (left), that interact with at least one virus (middle), and that interact with viral immunomodulators (viral proteins that target the host immune system) (right). Empirical p-values are shown. **(F)** A scaled heatmap showing: (1) values of response divergence (computed as in Fig 1C), (2) cell-to-cell variability (as in Fig 2A), (3) coding sequence divergence (dN/dS values, as in 3A), (4) gene age (as in 3C – younger genes have darker colours), (5) number of cellular interactions (as in 3D) and (6) number of host-virus interactions (as in 3E), for example genes from three functional groups: (I) cytokines, chemokines and their receptors, (II) transcription factors, (III) kinases and phosphates. Values are shown in a normalized scale between 0-100, so that in each column the gene with the highest value is given a score of 100 and the gene with the lowest value is given a score of 0. Genes that have missing values are coloured in grey.

### Transcriptionally divergent genes have higher rates of gene gain and loss in the course of mammalian evolution

Next, we tested the relationship between a gene’s divergence in response and the rate at which the gene’s family has expanded and contracted in the course of vertebrate evolution. Rapid gene duplication and gene loss have been observed in several important immune genes^36-41^. This has been hypothesized to be a result of pathogen-driven pressure^42,43^.

We used the statistics provided in ENSEMBL, where the relative rate of gene duplication and loss across vertebrates is assessed using a model based on global birth-and-death rates^44^. We found that transcriptionally divergent dsRNA-response genes have undergone faster rates of gene gain and loss (Fig 4B).

Higher rates of gene gain and loss in a gene family imply that at least some genes in these gene families are evolutionarily younger. We thus investigated the evolutionary age of the set of dsRNA-response genes using eleven different combinations of age estimation parsimonies and vertebrate gene families obtained from ProteinHistorian^45^. Indeed, genes that diverge more rapidly in their dsRNA-response are evolutionarily younger (Figs 4C Fig S15A). It is important to note that gene age estimations are not confounded by coding sequence evolution (Fig S15B). Thus, the two evolutionary mechanisms we studied - gene gain and loss and coding sequence evolution - are independent.

Previous reports suggest that younger genes tend to have fewer protein-protein interactions within cells^46^. We indeed observe that rapidly diverging genes (in both expression and protein sequence) tend to have fewer protein-protein interactions (obtained from STRING^47^). This agrees with the above finding on gene age (Fig 4D).

Together, these results suggest that transcriptionally divergent dsRNA-response genes evolve rapidly through various mechanisms, including fast coding sequence evolution and higher rates of gene loss and duplication events.

### Viruses evolve to preferentially interact with conserved innate immune genes

Pathogens are a major selective pressure in mammalian evolution^5,6^, and are thought to drive the evolution of immune genes. We thus investigated the relationship between transcriptional divergence and interactions with viral proteins by compiling a dataset of known host-virus interactions^5,48,49^. Interestingly, dsRNA-response genes with no known viral interactions have higher response divergence than genes with viral interactions (Fig 4E). Furthermore, the subset of dsRNA-response genes that are targeted by viral immunomodulators^50^ - viral proteins that subvert the host immune system - is even more conserved than the overall set of genes that interact with any viral protein (Fig 4E). This suggests that viruses have evolved to interact with innate immune proteins that are relatively conserved. Presumably, these conserved host genes cannot evolve away from these viral interactions^51^.

Our results are summarized in Fig 4F, which shows how representative dsRNA-response genes differ in both the evolutionary and regulatory characteristics we have studied. Cytokines, chemokines and their receptors evolve rapidly in all evolutionary modes and have higher transcriptional variability across cells. In contrast, dsRNA-induced genes that are involved in immune response regulation, such as kinases and transcription factors, are more conserved and less heterogeneous across cells. These genes have higher numbers of cellular protein-protein interactions, suggesting higher constraints imposed on their evolution. This latter group of conserved genes is more often targeted by viruses, revealing a relationship between host-pathogen dynamics and the evolutionary architecture of the innate immune response.

## Discussion

In this work, we have taken an integrative approach to chart the evolutionary architecture of the innate immune response. In particular, we have focused on the relationship between a gene’s cell-to-cell variability and its expression divergence between species. To dissect this relationship, we combined bulk and single-cell RNA-sequencing approaches with ChIP-sequencing experiments and molecular evolutionary analyses. We make this data available in an interactive web resource at: https://teichlab.sanger.ac.uk/ImmunityEvolution.

We find that rapidly diverging genes are also highly variable between cells. Both of these characteristics are associated with a particular promoter architecture. Moreover, genes with rapid expression divergence also evolve more rapidly in other evolutionary mechanisms: they are more divergent at the level of protein sequence, and undergo rapid gene gain and loss events.

Importantly, we find that genes involved in different functions in the innate immune response diverge at different rates. Genes related to response regulation are more conserved, while cytokines and chemokines diverge rapidly. Many cytokines and chemokines have high cell-to-cell variability, and are expressed in relatively few cells.

One of the central findings of our work is the observation that genes rapidly diverging between species exhibit higher levels of variability in their transcription across individual cells. Both characteristics - increased transcriptional divergence and heterogeneity across cells - are associated with a similar promoter architecture: genes whose promoters are enriched with TATA-boxes and depleted of CGIs, tend to be rapidly evolving and more variable between cells. This agrees with previous work in yeast, where CGI promoters were linked to low cell-to-cell variability and conservation in expression. TATA-box promoters were associated with genes with rapid transcriptional divergence between species and high variability in protein expression^31,52^. Interestingly, yeast genes with high transcriptional variability are often involved in stress responses and have a high dynamic range in their response^31^. This suggests intriguing analogies between the regulation of genes involved in stress response in yeast, and the regulation of the mammalian innate immune response, two systems that may have been subjected to accelerated evolution due to continuous changes in external stimuli.

Cytokines and chemokines display the highest transcriptional variability, being expressed in only a small subset of cells. This is observed across all stimulation conditions and species studied, and is in agreement with previous findings showing specific examples of cytokines expressed in a subset of responding cells^30^. We observe that cytokines tend to be co-expressed in the same cells. Moreover, they correlate in expression with genes that are known to be upstream regulators of the innate immune response, and this co-expression is conserved across species. This might indicate that the high cell-to-cell variability observed in cytokine expression is functionally important. Since prolonged or increased cytokine expression can result in tissue damage^53-55^, restricting cytokine production to only a few cells might assist in preventing pathological immune conditions such as autoimmunity. Therefore, expressing cytokines in just a few cells might be sufficient to activate the innate immune response in nearby cells and recruit immune cells, while retaining the ability to switch off the response in a controlled manner^9^.

In contrast to cytokines, we observe that genes involved in regulation of the immune response – such as transcription factors and kinases are less variable between cells, and are less diverging in transcriptional response along with other evolutionary mechanisms. These genes might have stronger functional and regulatory constraints, due to their roles in multiple contexts and pathways unrelated to innate immunity. These constraints limit their ability to evolve; an Achilles’ heel that can be used by pathogens to subvert the immune system. Indeed, we observe that viruses have evolved to preferentially interact with conserved factors in the innate immune response, possibly exploiting the constraints imposed on these genes that limit their ability to evolve away from viral interactions.

Due to these constraints, we suggest that a successful host strategy is manifested through cytokines and other secreted immune proteins. Not only do they act as an efficient means of alerting neighbouring and immune cells of pathogen presence, they also function as an important element in the evolutionary arms race. This stems from their ability to readily evolve with fewer constraints imposed by intracellular interactions or additional non-immune functions. This evolution manifests through multiple mechanisms: divergence in expression levels, changes in coding sequences and gene duplication and loss events.

Our work thus reveals the underlying architecture of the innate immune response, linking promoter structure and transcription with evolution, function and host-pathogen interactions. Our results suggest that rapid evolution and high levels of expression heterogeneity are prominent in genes with specific functions in the innate immune response. Those genes that are involved in regulation tend to be conserved, whereas cytokines diverge rapidly and exhibit high variability between cells. These characteristics may enable cytokines to induce a rapid, yet controlled, response across the tissue and to avoid long-lasting and potentially damaging effects.

## Acknowledgements

We would like to thank Nils Eling, Matteo Fumagalli, Yoav Gilad, Muzlifah Haniffa, Orly Laufman, Amir Marcovitz, John Marioni, Matthieu Muffato, Duncan Odom, Oliver Stegle, Adi Stern, Mike Stubbington and Michelle Ward for helpful discussions and advice throughout the project; Lira Mamanova and Sanger support teams for technical assistance, and members of the Teichmann lab for support in various stages. This project was supported by ERC grants (ThDEFINE and ThSWITCH) and an EU FET-OPEN grant (MRG-GRAMMAR). T.H was supported by a Human Frontier Science Program Long-Term Fellowship and by an EMBO Long-Term fellowship. T.G. by the European Union’s H2020 research and innovation programme “ENLIGHT-TEN” under the Marie Sklodowska-Curie grant agreement 675395.

## Materials and Methods

### Ethics

This project has been approved by the Wellcome Trust Sanger Institute Animal Welfare and Ethical Review Body (AWERB). Human samples have been obtained from open-access lines from the Human Induced Pluripotent Stem Cells Initiative (HipSci)^1^. Non-human primate skin samples were obtained from animals assigned to unrelated non-infectious studies, provided by Public Health England’s National Infection Service (PHE) in accordance with Home Office (UK) guidelines and approved by the Public Health England Ethical Review Committee under an appropriate UK Home Office project license; and by the Biomedical Primate Research Centre in the Netherlands (BPRC) in accordance with the Dutch Law on Animal Experimentation, following EC Directive 2010/63/EU on the protection of animals used for scientific purposes.

### Data access

Our results can be visualized in an online webserver - https://teichlab.sanger.ac.uk/ImmunityEvolution - where users can visualize histone marks across species and individuals, fold changes in response to dsRNA and IFNB across species from bulk transcriptomics, as well as the gene expression values in individual cells across species and stimulation time course.

Sequencing data is deposited in ArrayExpress and would become available upon publication.

### Experimental methods

#### Tissue culture

We cultured primary dermal fibroblasts from low passage cells (below 10) that originated from sexually mature females from four different species (human (European ancestry), rhesus macaque, black 6 mouse and brown Norway rat). All skin samples originated from shoulders and were extracted and grown in a similar manner. Stimulation experiments and library preparations were done in identical conditions across all species and for all genomics techniques. For each species and for each genomics technique we used several biological replicates. Details on the numbers of individuals used in each technique are detailed in each technique’s section and in Table S1.

Human cells were obtained from the Hipsci project (http://www.hipsci.org/ ^1^). Non-human primate cells were extracted from skin tissues that were incubated for 2hrs with 0.5% collagenase-A (Roche; 11088815001) after mechanical processing, and then filtered through 100µm strainers before being plated and passaged prior to cryo-banking. Rodent cells were obtained from Cellbiologics where they were extracted using a similar enzymatic extraction protocol using skin tissues from the same regions of the body used in primates.

Prior to stimulation, cells were thawed and grown for 3 days in ATCC fibroblast growth medium (Fibroblast Basal Medium (ATCC, ATCC-PCS-201-030) with Fibroblast Growth Kit-Low serum (ATCC, PCS-201-041) (supplemented with Primocin (Invivogen, ant-pm-1) and Pen/Strep (Life Technologies, Cat. Code: 15140122))– a controlled medium that has proven to provide good growing conditions for fibroblasts from all species, having slightly less than 24h doubling times. ∼18hrs before stimulation, cells were trypsinized, counted and 100,000 cells were seeded into 6-well plates, leading to ∼60-70% confluence in the beginning of the stimulation experiment. Cells were stimulated with either: (1) 1ug/mL High-Molecular Weight poly I:C (Invivogen, Cat. Code: tlrl-pic) transfected with 2uL/mL Lipofectamin 2,000 (ThermoFisher, Cat Number 11668027) (these conditions lead to ∼100% transfection efficiency across species within 2 hours post-transfection); (2) mock transfected with Lipofectamin 2,000; (3) stimulated with 1,000 IU of IFNB for 8 hours (human IFNB - 11410-2 (for human and macaque cells); rat IFNB - 13400-1; mouse IFNB – 12401-1; all IFNs were obtained from PBL, and had activity units based on similar virological assays); or (4) left untreated.

Additional samples in human and mouse were stimulated with 1,000 IU of Cross-mammalian IFN (CMI, or Universal Type I IFN Alpha, PBL, Cat Number 11200-1). The latter stimulation was done to assess the effects of species-specific and batch-specific IFNB: Using cross-mammalian IFN allows stimulating cells from different species with the same IFN, thus allowing to infer the species-specific response to IFN. The concentrations and time points used were chosen following 24-hour time-course calibration experiments and after testing different concentrations.

Human fibroblasts were also cultured in an alternative experimental condition to generate a larger dataset of single cells and enable confirmation of results across experimental settings (this batch is referred to as the third replicated in the single-cell analysis). For this, cells were grown in DMEM, high glucose, GlutaMAX Supplement, pyruvate (Life Technologies, Cat code: 10569-010), 10% Fetal Bovine Serum (Lab Tech, Cat code: FB-1001) and Penicillin – Streptomycin (Life Technologies, Cat code: 15140122). Cells were stimulated for 2 and 6 hours with 0.5ug/mL High-Molecular Weight poly I:C (Invivogen, Cat. Code: tlrl-pic) transfected with 1uL/mL Lipofectamin 2,000 (ThermoFisher, Cat Number 11668027); or left untreated, before trypsinization and processing.

### Bulk RNA sequencing

#### Library preparation and sequencing

For bulk transcriptomics analysis, individuals from different species were grown in parallel and stimulated with dsRNA, IFNB (and cross-mammalian IFN) and with their respective controls. In total, samples from 6 humans, 6 macaques, 3 mice and 3 rats were used (we used 3 samples for rodents due to their lower genetic variability). Total RNA was extracted using the RNeasy Plus Mini kit(Qiagen, Cat Number 74136), using QIAcube (Qiagen). RNA was then measured using a Bioanalyzer 2100 (Agilent Technologies), and samples with RIN<9 were excluded from further analysis.

Libraries were produced using the Kapa Stranded mRNA-seq Kit (Kapa Biosystems, Kit code: KK8421). The Kapa library construction protocol was modified for automated library preparation by Bravo (Agilent Technologies). cDNA was amplified in 13 PCR cycles, and purified by Ampure XP beads (Beckman Coulter, Cat Number A63882) (1.8x volume) using Zephyr (Perkin Elmer). Prior to sequencing, libraries were quantified using MiSeq (25bp PE). This data was used for pooling libraries in equimolar amounts by Coulter NX (Span-8, Beckman). Pooled samples were sequenced on an Illumina HiSeq 2500 instrument, using paired-end 125-bp reads.

### ChIP sequencing

#### Library preparation and sequencing

Samples from three individuals from each of the four species were grown and stimulated (with poly I:C or left untreated, as described above) in parallel to samples collected for bulk RNA-seq. Following stimulation, sample were crosslinked in 1% HCHO (prepared in 1X DPBS) at room temperature for 10 minutes, and HCHO was quenched by the addition of glycine at a final concentration of 0.125M. Cells were pelleted at 4°C at 2000 x g, washed with ice-cold 1X DPBS twice, and snapped frozen in liquid nitrogen. The cell pellets were stored in -80°C until further stages were performed. ChIPmentation was performed according to version 1.0 of the published protocol ^2^ with some modifications at the ChIP stage:

Cell pellets were lysed in 300ul ChIP Lysis Buffer I (50mM HEPES.KOH, pH 7.5, 140mM NaCl, 1mM EDTA, pH 8.0, 10% Glycerol, 0.25% NP-40, 0.5% Triton X-100) on ice for 10 minutes. Cells were then pelleted at 4°C at 2000 x g for 5 minutes, and washed by 300ul ChIP Lysis Buffer II (10 mM Tris.Cl, pH 8.0, 200 mM NaCl, 1mM EDTA, pH 8.0, 0.5mM EGTA, pH 8.0), and pelleted again at 4°C at 2000 x g for 5 minutes. Nuclei were resuspended in 300ul ChIP Lysis Buffer III (10mM Tris.Cl, pH 8.0, 100mM NaCl, 1mM EDTA, 0.5mM EGTA, 0.1% Sodium Deoxycholate, 0.5% N-Lauryolsarcosine). Chromatin was sonicated using Bioruptor Pico (Diagenode) with 10 cycles of 30 seconds ON/30 seconds OFF. 30ul 10% Triton X-100 were added into each sonicated chromatin, and insoluble chromatin was pelleted at 16,100 x g at 4°C for 10 minutes. 1ul supernatant was taken as an input control. The rest of the supernatant was incubated with 10ul Protein A Dynabeads (Invitrogen) pre-bound with 1ug anti-H3K4me3 (Diagenode Cat Number C15410003) or 1ug anti-H3K27ac (Diagenode, Cat Number C15410196) in a rotating platform in a cold room overnight. Each immunoprecipitation (IP) was washed with 500ul RIPA Buffer (50mM HEPES.KOH, pH 7.5, 500mM LiCl, 1mM EDTA, 1% NP-40, 0.7% Sodium Deoxycholate, check components) for 3 times. Each IP was then washed with 500ul 10mM Tris, pH 8.0 twice, and resuspended in 30ul tagmentation reaction mix (10mM Tris.Cl, pH 8.0, 5mM Mg2Cl, 1ul Tn5 transposase (Nextera)). The tagmentation reaction was put on a thermomixer at 37°C for 10 minutes at 800 rpm shaking. Following tagmentation, each IP was washed sequentially with 500ul RIPA Buffer twice, and 1X TE NaCl (10mM Tris.Cl, pH 8.0, 1mM EDTA, pH 8.0, 50mM NaCl) once. Elution and reverse-crosslinking were done by resuspending the beads with 100ul ChIP Elution Buffer (50mM Tris.Cl, pH 8.0, 10mM EDTA, pH 8.0, 1% SDS) on a thermomixer at 65°C overnight, 1,400 rpm. DNA was purified by MinElute PCR Purification Kit (QIAGEN, Cat Number 28004) and eluted in 12.5ul Buffer EB (QIAGEN, Cat Number 28004), which yielded ∼10ul ChIPed DNA.

The library preparation reactions contained the following reagents: 10ul purified DNA (from the above procedure), 2.5ul PCR Primer Cocktails (Nextera kit, Illumina, Cat Number FC-121-1030), 2.5ul N5xx (Nextera index kit, Illumina Cat Number FC-121-1012), 2.5ul N7xx (Nextera index kit, Illumina, Cat Number FC-121-1012), 7.5ul NPM PCR Master Mix (Nextera kit, Illumina, Cat Number FC-121-1030) The PCR cycles were as follows: 72°C, 5 mins; 98°C, 2 mins; 98°C, 10 secs, 63°C, 30 secs, 72°C, 20 secs] X 12; 10°C hold.

The amplified libraries were purified by double AmpureXP beads purification: first with 0.5X bead ratio, keep supernatant, second with 1.4X bead ratio, keep bound DNA. Elution was done in 20ul Buffer EB (QIAGEN).

1ul of library was run on an Bioanalyzer (Agilent Technologies) to verify normal size distribution. Pooled samples were sequenced on an Illumina HiSeq 2000 instrument, using paired-end 75-bp

### Single-Cell RNA sequencing

#### Flow cytometry

For scRNA-seq, we performed two different biological replicates, with each replicate having one individual from each of the four studied species. A time course of dsRNA stimulation of 0,2,4, and 8hrs was used in one replicate, and a smaller replicate included 0,4,8hrs. All cells from the same biological replicate were sorted in identical conditions and into identical plates. Poly I:C transfection was done as described above. In the case of sorting with IFNLUX, we used a rhodamine-labelled poly I:C, to collect index data on poly I:C transfection.

Cells were sorted with either Becton Dickinson INFLUX (first replicate) or Beckman Coulter MoFlo XDP (second replicate) into wells containing 2uL of Lysis Buffer (1:20 solution of RNase Inhibitor (Clontech, Cat Number 2313A) in 0.2% v/v Triton X-100 (Sigma-Aldrich, Cat Number T9284), spun down and immediately frozen at -80°. Technical positive and negative controls of small bulk with 10 cells or with empty wells respectively, were used to assess the quality of the processed cells. Cells from different conditions (time points or species) were sorted based on a Latin square design to minimize biases due to specific locations on the plate. These plates were later processed together to minimize batch effects.

When sorting with MoFlo (smaller biological replicate), a pressure of 15psi was used with a 150µm nozzle, using the ‘Single’ sort purity mode. Dead or late-apoptosis cells were excluded using Propidium Iodide at 1µg/ml (Sigma, Cat Number P4170) and single cells were selected using FSC W vs FSC H.

When sorting with INFLUX (larger biological replicate), a pressure of 3psi was used with a 200µm nozzle, with the “single” sort mode. Index data was collected and the median fluorescence intensity (MFI) of rhodamine-labelled poly I:C (InvivoGen, tlrl-picr), DAPI, forward and side scatter was retained. Dead or late-apoptosis cells were excluded using 100ng/ml DAPI (4’,6-diamidino-2-phenylindole) (Sigma, Cat Number D9542). DAPI was detected using the 355nm laser (50mW) using a 460/50nm bandpass filter. Rhodamine was detected using the 561nm laser (50mW), using a 585/29nm bandpass filter. Single cells were collected using the FSC W vs FSC H. In addition to the two described replicates that included cells from all four species, human cells from 3 individuals were sorted in a third experiment (in parallel to cells from the same individuals and conditions being processed using microfluidic droplet cell capture). Cells were sorted with INFLUX as described above.

#### Library preparation from full-length RNA from single cells and sequencing

Sorted plates were processed according to the Smart-seq2 protocol^3^: Oligo-dT primer (IDT), dNTPs (ThermoFisher, Cat Number 10319879) and ERCC RNA Spike-In Mix (1:25,000,000 final dilution, Ambion, Cat Number 4456740) were added to each well, and Reverse Transcription (using 50U SmartScribe, Clontech Cat Number 639538) and PCR were performed following the original protocol with 25 PCR cycles. cDNA libraries were prepared using Nextera XT DNA Sample Preparation Kit (Illumina, Cat Number FC-131-1096), according to the protocol supplied by Fluidigm (PN 100-5950 B1). Quality Checks on cDNA were done using a Bioanalyser 2100 (Agilent Technologies). Libraries from 96 single cells were quantified using the LightCycler 480 (Roche), pooled and purified using AMPure XP beads (Beckman Coulter) with Hamilton 384 head robot (Hamilton Robotics). Pooled samples were sequenced on an Illumina HiSeq 2500 instrument, using paired-end 125-bp reads.

#### Library preparation from single cells using microfluidic droplet cell capture

Cells from three human individuals were grown, stimulated with dsRNA for 6hrs or left unstimulated, collected by trypsinization and processed in a 10X Chromium machine (10X Genomics). Following multiplexing, libraries were prepared according to the manufacturer’s protocol. Pooled samples were sequenced on an Illumina HiSeq 2500 instrument, using paired-end 75-bp reads. We used this approach as a second experimental technique to prepare libraries from single cells, to confirm our results regarding cell-to-cell variability, obtained from the primary method we use (Smart-seq2). This method is more highly parallel in cell capture but less sensitive than our primary plate-based method (Smart-seq2) as a single cell RNA-seq technology^4^.

## Computational methods

### Bulk RNA transcriptomics

#### Read mapping to annotated transcriptome

Adaptor sequences and low-quality score bases were first trimmed using Trim Galore (version 0.4.1) (with the parameters “--paired --quality 20 --length 20 –e0.1 --adapter AGATCGGAAGAGC”). Trimmed reads were mapped and gene expression was quantified using Salmon (version 0.6.0) ^5^ with the following command: “salmon quant -i [index_file_directory] -l ISR -p 8 --biasCorrect --sensitive --extraSensitive -o [output_directory] -1 -g [ENSEMBL_transcript_to_gene_file] --useFSPD --numBootstraps 100”. Each sample was mapped to its respective species’ annotated transcriptome (downloaded from ENSEMBL, version 84: GRCh38 for human, MMUL_1 for macaque, GRCm38 for mouse, Rnor_6.0 for rat). We only included the set of coding genes (*.cdna.all.fa files). In the case of human, we removed annotated secondary haplotypes of coding genes. We did not include long non-coding RNAs since coding genes are the focus of this study and since current annotations of long non-coding RNAs significantly differ in numbers and quality between species.

#### Quantifying differential gene expression in response to dsRNA or IFNB

To quantify differential gene expression between treatment and control for each species and for each treatment separately, we used edgeR (version 3.12.1)^6^ using the rounded estimated counts returned by Salmon in the previous stage. This was done only for genes that had a significant level of expression in at least one of the four species (TPM>3 in at least N-1 libraries, where N is the number of different individuals we have for this species with libraries that passed quality control). Significance testing was done using the exact test from edgeR, and p-values were corrected using FDR.

### Conservation and divergence in response

#### Fold change-based phylogeny

We compared the overall change in response to treatment (dsRNA and to IFNB) between pair of species, by computing the Spearman correlation of the fold change in response treatment across all one-to-one orthologs that were expressed in at least one species (Figs S1A-H). Fold change was calculated with edgeR, as explained above.

We constructed a tree based on gene’s changes in expression in response to dsRNA and to IFNB using 9,835 expressed genes that had one-to-one orthologs across all four species and were expressed in at least one species in at least one condition (Fig S1I). We used hierarchical clustering, using the command hclust from the stats R package (version 3.3.2), with the distance between samples computed as 1-*ρ*, where *ρ* is the pairwise Spearman correlation between each pair of species mentioned above (and thus, a greater similarity, reflected in a higher correlation, results in a smaller distance) and “average” as the clustering method. We note that when repeating this procedure with gene’s p-values we obtained a similar phylogeny.

#### Quantifying gene expression divergence using Orenstein-Uhlenbeck models

The Ornstein-Uhlenbeck (OU) process was suggested as an appropriate model to study the evolution of gene expression^7^ and was previously used to model the evolution of gene expression in various systems, such as in mammalian tissues^8^ and in fly embryonic development^9^.

OU models incorporate selection and drift, and accordingly include two terms reflecting stochastic and deterministic forces. The model describes the change of a quantitative character *X* over a small increment of time *t* as: *dX(t)* = α[θ − *X(t)*]*dt* + *σdB(t)*, where the first term describes the deterministic force of selection, and the second term is the stochastic Brownian motion representing genetic drift.’ *α* is the strength of selection, *σ* is the strength of neutral drift and (is the trait’s optimum.

We used the ouch R package (version 2.9-2) to fit OU and Brownian motion models to our data of fold change in gene expression across species. We fitted 8 different models to the species-specific fold change values for each expressed gene. We used a phylogenetic tree, which was constructed using the concatenated nucleic acid sequences of all the expressed genes in each expression program (dsRNA or IFNB) that had one-to-one orthologs across the four species (MSA was created using MUSCLE, version 3.7^10^), and tree was reconstructed using PhyML, version 3.0^11^). The models were a pure Brownian motion model, reflecting neutral evolution, and 7 Hansen models with differing number of selective optima. Coefficients values for selection and drift (*α* and *σ*, respectively) were then retrieved from the model that best fitted the data, based on the Akaike Information Criterion. A measure of divergence in gene expression response was calculated from these values as: 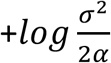 ^7^. For simplicity, throughout the manuscript we refer to this measure as “response divergence”.

We note that here we did not attempt to develop an extensive model, but rather to use an existing model that captures changes in expression between species while taking into account their known evolutionary relationships. Importantly, when we compare the divergence values obtained with the OU model with divergence values obtained from fitting the data with linear models (see below) – we get a significant agreement between the two approaches (see Fig S1A-F).

For various analyses where we compare response divergence with another characteristics (e.g. – coding sequence evolution, active chromatin marks divergence, cell-to-cell variability, etc.), the 955 dsRNA-response genes (genes that are differentially expressed in humans, with a corrected p-value<0.01, based on edgeR exact test) were split into three groups, based on their level of response divergence across species: (1) highly divergent genes (the top 25% genes with the highest divergence values), (2) lowly divergent genes (the bottom 25%), and (3) genes with medium level of divergence (the middle 50%).

#### Quantifying gene expression divergence using linear models in bulk transcriptomics and in single cell time course transcriptomics

To quantify gene expression divergence using a different approach to the above-mentioned OU models, we fitted a linear model to the bulk expression data of all the expressed orthologous genes in response to treatment (dsRNA or IFNB) in each pair of species. We used the glmFit command in the edgeR R package (version 3.12.1), to fit a linear equation that incorporates species and treatment, as follows: *y = aT + bS + cT + e*, where T is the treatment effect, S is the species effect, T:S is the interaction between treatment and species, and = is the residual error. We then extracted the fold-change and p-value for the interaction term (c), and used them as a proxy for divergence in expression between species in response to treatment. When comparing the two models (OU and linear models), we observe a significant agreement between the two models (see Fig S3). Thus, the group of genes that are divergent in response across species is robust to the computational approach we use to detect them.

#### Comparison of response divergence between different functional groups

To compare the divergence rates between sets of dsRNA-response genes that have different functions in the innate immune response, we split these 955 genes into the following functional groups (all groups are mutually-exclusive, and any gene that belongs to two groups was excluded from the latter group):

We first grouped genes by annotated molecular functions: Viral sensors (genes that belong to one of the GO categories: GO:0003725 - double-stranded RNA binding, GO:0009597 - detection of virus; GO:0038187 - pattern recognition receptor activity), Cytokine & chemokine receptors (GO:0004896 - cytokine receptor activity; GO:0004950 - chemokine receptor activity), Cytokines & chemokines (GO:0005125 cytokine activity; GO:0008009 chemokine activity), Transcription factors (taken from the Animal Transcription Factor DataBase (version 2.0) ^12^), Chromatin modulators (GO:0016568 - chromatin modification, GO:0006338 - chromatin remodelling, GO:0003682 - chromatin binding, GO:0042393-histone binding), Kinases and phosphatases (GO:0004672 - protein kinase activity; GO:0004721 - phosphoprotein phosphatase activity), Ligases and deubiquitinases (GO:0016579 protein deubiquitination; GO:0004842 - ubiquitin-protein transferase activity GO:0016874 ligase activity), Other enzymes (mostly involved in metabolism rather than regulation - GO:0003824 - catalytic activity). The divergence response values of these functional subsets were compared to the entire group of dsRNA-response genes.

Next, we grouped genes by biological processes that are known to be important in the innate immune response: Cellular defence (GO:0051607 - defence response to virus), Inflammation (GO:0006954 - inflammatory response), Apoptosis (GO:0006915 - apoptotic process), and Regulation (the annotations of regulation that are specific to the dsRNA-IFN pathway include only few genes. We thus merged the genes that are annotated as transcription factors, chromatin modulators, kinases and phosphates or ligases and deubiquitinases, since all these group include numerous genes that are known to regulate the innate immune response.)

Gene lists belonging to the mentioned GO annotations were downloaded using QuickGo ^13^. The distribution of response divergence values for each of the functional groups was compared with the distribution of response divergence of the entire set of dsRNA-response genes. Plots of functional groups of genes in the dsRNA response are shown in Fig 2 (where genes are categorized by molecular functions (2C) or by biological process (2D)). Analogous plots for functional groups in the IFNB response (with 841 IFNB-response genes) are shown in Fig S5.

### Conservation of active chromatin marks analysis

#### Alignment and peak calling of ChIP-seq reads

ChIP-seq reads were trimmed using trim_galore (version 0.4.1) with “--paired --trim1 --nextera” flags. The trimmed reads were aligned to the corresponding reference genome (hg38 for human, rheMac2 for macaque, mm10 for mouse, rn6 for rat – all these genomes correspond to the transcriptomes used for RNA-seq mapping) from the UCSC Genome Browser ^14^ using bowtie2 (version 2.2.3) with default settings ^15^. In all four species we removed the Y chromosome. In the case of human, we also removed all alternative haplotype chromosomes. Following alignment, low-confident mapping and improperly-paired reads were removed by samtools ^16^ with “-q 30 –f 2” flags.

Enriched regions (peaks) were called using MACS2 (v 2.1.1)^17^ with a corrected p-value cutoff of 0.01 with “-f BAMPE -q 0.01 -B --SPMR” flags, using input DNA as control. The genome sizes (the argument for “-g” flag) used were “hs” for human, “mm” for mouse, 3.0e9 for macaque and 2.5e9 for rat. Peaks were considered reproducible when they were identified in at least two of the three biological replicates and overlapped by at least 50% of their length (non-reproducible peaks were excluded from subsequent analyses). Reproducible peaks were then merged to create consensus peaks from overlapping regions of peaks from the three replicates, by using mergeBed from the bedtools suite^18^.

#### Gene assignment and conservation of active promoters and enhancers

We subsequently linked human peaks with the genes they might be regulating as follows: H3K4me3 consensus peak was considered the promoter region of a given gene if its centre was between 2kb upstream and 500bp downstream of the annotated TSS of the most abundantly-expressed transcript of that gene. Similarly, H3K27ac was considered the enhancer region of a given gene if its centre was in a distance above 1kb and below 1Mb, and there was no overlap (of 1bp or more) with any H3K4me3 peak.

In each case where, based on the distance criteria, more than a single peak was linked to a gene (or more than a single gene was linked to a peak), we took only the closest peak-gene pair (ensuring that each peak will have up to one gene and vice versa).

To compare active promoters and enhancers between species, we excluded any human peak that could not be uniquely mapped to the respective region in the other species. This was done by looking for syntenic regions of human peaks in the other three species by using liftOver ^19^, and removing peaks that have either unmapped regions or more than one mapped region in the compared species. We considered syntenic regions with at least 10% sequence similarity between the species (minMatch=0.1), with a minimal length (minSizeQ and minSizeT) corresponding to the length of the shortest peak (128bp in H3K4 and 142bp in H3K27).

We defined an active human promoter or enhancer as conserved if a peak was identified in the corresponding region of the other species (we repeated this analysis by comparing human with each of the other three species, separately). We compared the occurrence of conserved promoters and enhancers in genes that are highly divergent in response to dsRNA with lowly divergent genes, and used Fisher exact test to determine statistical significance of the observed differences between highly and lowly divergent genes. Results are shown in Figure S4.

### Promoter sequence analysis

#### Promoter sequence evolution analysis

PhyloP7 values were used to assess promoter sequence conservation^20^. Sequence conservation quantification was done by taking the estimated nucleotide substitution rate for each nucleotide along the promoter sequence (500bp upstream of the TSS). When several annotated transcripts exist, the TSS of the most abundantly expressed transcript was used. The substitution rate values from all genes were aligned, based on their TSS position, and a mean for each of the 500 positions was calculated separately for the group of genes with high, medium and low response divergence. Two-sample Kolmogorov-Smirnov test was used to compare the paired distribution of rates between the mean of the highly divergent and the mean of the lowly divergent sets of genes. To plot the mean values of the three sets of divergent genes, geom_smooth function from the ggplot2 R package was used with default parameters (with loess as the smoothing method) (see Fig 2A).

#### Promoter elements analysis

Human CpG island (CGI) annotations were downloaded from the UCSC genome table browser (hg38), and CGI genes were defined as those with a CGI overlapping their core promoter (300bp upstream of the TSS reference position, and 100bp downstream of it, as suggested previously^21^). Genes were defined as having a TATA box if they had a significant match to the Jaspar TATA box matrix (MA0108.1) in the 100bp upstream of their TSS by FIMO^22^ with default settings (we note that only 28 out of 955 dsRNA-response genes had a matching TATA-box motif in this region).

### Cell-to-cell transcriptional variability analysis

#### Read mapping and quality control of single-cell RNA-SEQ (full-length RNA)

Gene expression was quantified in a similar manner to the description of bulk transcriptomics libraries, with the addition of ERCCs to the species’ reference transcriptomes. Low-quality cells were filtered following quality control criteria (cells with at least 250,000 mapped reads, with at least 2,000 expressed genes with TPM>3, with ERCC<10% and MT<40%), resulting in 728 cells from one biological replicate (with a dsRNA stimulation time course of 0,2,4,8 hours) and 240 cells from the second biological replicate (with a time course of 0,4,8 hours). A third replicate of human cells resulted in 590 high quality cells. Results throughout the manuscript relate to the larger cross-species biological replicate, unless otherwise specified.

#### Quantifying gene expression in microfluidic droplet cell capture data

Gene expression was quantified using 10X Genomics’ Cell Ranger pipeline for alignment, de-duplication and filtering (version 1.2.0) (https://support.10xgenomics.com/single-cell/software/pipelines/latest/what-is-cell-ranger). Low-quality cells were removed, so that only cells with at least 1,000 expressed genes were included, resulting in a total of 4,468 high-quality cells.

#### Cell-to-cell variability analysis

To quantify gene’s biological cell-to-cell variability we have applied two different and complementary computational approaches:

(1) DM (Distance to Median) - an established method, which calculates the cell-to-cell variability in gene expression, while accounting for the confounding factors of gene expression level^23^. This is done by first filtering out genes that are lowly expressed: For Smart-seq2 data we only included genes that have an average expression of more than 10 reads. For microfluidic droplet cell capture data, we used genes that had been detected with a UMI of at least 1 in at least 20 cells. This filtering procedure is done to avoid genes that display higher levels of technical variability between samples. Secondly, to account for gene expression level, the observed cell-to-cell variability of each gene is compared with its expected variability, based on its mean expression across all samples and in comparison, with a group of genes with similar levels of mean expression. This distance from mean value (DM), is also corrected by gene length, yielding a value of variability that can be compared across genes regardless of their length and mean expression values^24^.

Results from DM analysis are shown in Figure 3A-C (human cells, stimulated with poly I:C for 4 hours), Figure 4D (human cells, different stimulation time points), Figure S7 (all four species, all time points), and Figure S8 (human cells processed with microfluidic droplet capture). Analysis was repeated with the second cross-species replicate and gave similar results (data not shown). In Figures 3D and S12 we show cell-to-cell variability values in a comparative manner across conditions. Specifically for these plots, we used a set of genes that have at least 5 reads in each of the stimulated conditions (ensuring a comparable set of expressed genes across all conditions). We used this set of mutual genes, to compute DM for each of the stimulated time points.

(2) As a second approach, we used BASiCS^25^ – a method that is based on a Bayesian algorithm developed specifically to identify truly biological variation in gene expression in single-cell transcriptomics data. To regress out variation that stems from technical well-to-well variation, it uses the variation in externally spiked-in RNA molecules^26^ to infer what is the likely level of technical noise, based on each gene’s variation and overall expression.

We used BASiCS with defaults parameters, taking into account genes that were expressed in at least two cells (with TPM>1). We verified that the model’s parameters were converging prior to the 20,000 iterations “burn period”, and ran the model for a further 20,000 iterations using a thinning period of 10. For each gene, the parameter representing biological variability (*δ*) was extracted.

Results from BASiCS analysis are shown in Figure S9.

#### Analysis of cell states in the innate immune response

To look at cell response levels in depth, we used the third replicate of human cells (from which we have the largest numbers of cells and from which we have processed cells from two different library preparation methods (10X microfluidic droplet capture and Smart-seq2).

Previous studies using different systems have indicated that cells responding to pathogen-associated molecular patterns such as poly I:C or LPS, often display different levels of response, where some cells exhibit a higher level of response than other responding cells^27,28^.

To study the effects of these response levels on gene cell-to-cell variability, we used 590 human cells that were stimulated with poly I:C for 2 or 6 hours, or left unstimulated. We plotted the results of a Principal Component Analysis (PCA) on these cells, and observed that the first PC distinguishes a population of cells that are closer to the unstimulated cells (lowly responsive cells), from a population that is further away (highly responsive cells) (Fig S10A). As expected, the later stimulation time point (6 hours) has a higher fraction of highly responding cells than the earlier time point.

These two populations display different levels of expression of dsRNA-upregulated genes (Fig S10B). We define dsRNA-upregulated genes as the set of genes exclusively upregulated in the primary response, following dsRNA treatment (having a corrected p-value<0.01 in edgeR DE analysis), which are not stimulated by IFNB (having p-values>0.3 in edgeR DE analysis of IFNB. These two sets of responding cells also differ in the fraction of cells expressing IFNB (Fig S10C). We use these genes to define two clusters of cells (using hierarchical clustering, with ward.D2 as the method, and with Euclidian distance) – see Figure S10D. The left cluster constitutes the “lowly responding cells", which cluster together with the unstimulated cells, whereas the “highly responding cells” appear in the right cluster.

We repeated the cell-to-cell variability analysis using the DM method on each population (separately for each of the two populations at 2 hours and at 6 hours). Using these DM values, we plot the divergence of lowly-, medium- and highly-divergent genes (as in Fig 3A), separately for highly and lowly responsive cells in Fig S10E. Similar to the analysis done in Main Fig 3A (where we did not separate cells by response levels), highly diverging genes have higher cell-to-cell variability in each cell state and in each time point.

#### Cytokine co-expression analysis

For analysis of co-expression within the set of cytokines, we correlated the expression of these genes in our largest dataset - 1,922 human cells (Spearman correlation), which were stimulated with dsRNA for 6 hours and processed using microfluidic droplet cell capture (see Fig S14).

For correlation of cytokines with all genes expressed across stimulated cells and in each species, we calculated Spearman correlation of expression between pairs of genes in dsRNA-stimulated cells (dsRNA-transfected cells for 2, 4 and 8 hours) for each of the four species. For the 13 cytokines that were found to cluster in the human data (Fig S14), we define a set of ‘cytokine-correlated’ genes - those with a correlation of ρ > 0.3 - as genes that highly correlate with cytokine expression in response to dsRNA. We ranked these genes (based on how many cytokines each gene was found to be correlated with) and used GOrilla^29^ to find enriched GO terms (Table S3A).

Using the 13 cytokines described above, we repeated co-expression analysis in each of the other 3 species (macaque, rat and mouse), calculating correlation with all expressed genes. We then used the categorisation of ‘cytokine-correlated’ from human data (described above) to split genes into two categories: those which correlate with cytokines in human, and those which are ‘non-correlated’. Using this grouping, we compared the correlation values between the ‘cytokine-correlated’ and ‘non-correlated’ in each species using a Kolmogorov-Smirnov test. We observe that the ‘cytokine-correlated’ genes have higher correlation values in macaque, mouse and rat than ‘non-correlated’ genes (p=2.0x10-77 in macaque, p=3.1x10-36 in rat, p=3.0x10-25 in mouse).

For the chemokine CXCL10, we built a network (using CytoScape^30^) of genes that correlate with CXCL10 in dsRNA-stimulated human cells and in at least one more species. The full network consists of 34 genes and is shown in Fig S14. A subset of this network is shown in Fig 3E, where we only include cytokine and chemokines and genes known to be involved in upregulation of cytokines (having the GO terms: “positive regulation of innate immune response” or “type I interferon signalling pathway”).

### Analysis of different evolutionary modes

#### Coding sequence evolution analysis

The ratio dN/dS of non-synonymous to synonymous codon substitutions, of human genes across the mammalian clade was obtained from a previous study using orthologous genes from 29 mammals^31^. Distributions of dN/dS values were computed for each of the three groups of genes with low, medium and high divergence in response to dsRNA, and are plotted in Figs 4A. The distributions of dN/dS values in lowly and highly divergent genes were compared using Mann-Whitney test.

We also compared response divergence with dN/dS values estimated in two additional sources: (1) dN/dS values computed based on 24 mammalian genomes^32^, and (2) pairwise dN/dS values of human-mouse, human-rat and human-macaque taken from ENSEMBL. In all cases when using the different dN/dS estimates, we observe the same trend, where genes that are highly divergent in response to dsRNA have higher dN/dS values in comparison with lowly divergent genes (data not shown).

#### Rate of gene gain and loss analysis

The significance at which gene’s family has experienced higher rate of gene gain and loss in the course of vertebrate evolution, in comparison with other gene families, was retrieved from ENSEMBL^33^. The statistics provided by ENSEMBL is calculated based on the CAFE method^34^, which estimates the global birth and death rate of gene families and identifies gene families that have accelerated rates of gain and loss. Distributions of the p-values from this statistic were computed for each of the three groups of genes with low, medium and high divergence in response to dsRNA and are plotted as the negative logarithm values in Fig 4B. The distributions of lowly and highly diverging genes were compared using Mann-Whitney test.

#### Gene age analysis

Gene age estimations were obtained from ProteinHistorian^35^. To ensure that the results are not biased by a particular method of ancestral protein family reconstruction or by specific gene family assignments, we used eleven different estimates for mammalian genes (combining five different databases of protein families with two different reconstruction algorithms for age estimation, as well as an estimate from the phylostratigraphic approach). For each gene, age is defined with respect to the species tree, where a gene’s age corresponds to the branch in which its family is estimated to have appeared (thus, larger numbers indicate evolutionarily older genes).

The age distributions of lowly and highly diverging genes were compared using a Mann-Whitney test. Data for gene age in comparison with divergence in response to dsRNA is shown in Fig 4C (using Panther7 phylogeny and Wagner reconstruction algorithm) and in Fig S16A (for all 11 combinations of gene family assignments and ancestral family reconstructions).

In addition, we tested whether results regarding gene age estimation are confounded by the ability to detect orthologs, which can be affected by the rate in which a gene’s coding sequence has diverged. For this, plotted in Fig S15B, we separated the 955 dsRNA-response genes based on their coding sequence divergence, into four quartiles. In each quartile, we plot the estimation of gene age for the genes that are highly-, medium-, or lowly-diverging in response to dsRNA. Gene age estimations are shown using four methods of gene estimation. In each case, we observe that the trend of highly-divergent genes to be younger than lowly-divergent genes is maintained.

#### Cellular protein-protein interactions analysis

Data on number of experimentally-validated protein-protein interactions (PPIs) for human genes was obtained from STRING (version 10)^36^. Distributions of PPIs for genes with low, medium and high divergence in response to dsRNA are plotted in Fig 4D. The distributions of lowly and highly diverging genes were compared using Mann-Whitney test.

#### Host-virus interactions analysis

Data on host-virus protein-protein interactions was downloaded from the VirusMentha database^37^, and combined with two additional papers that have annotated host-virus protein-protein interactions^32,38^. We split the 955 dsRNA-response genes into genes with known viral interactions (genes whose protein products were reported to interact with at least one viral protein), and genes with no known viral interactions: “Viral Interactors” and “No viral Interactions” respectively, in Figure 4E. In addition, we define a subset of genes within the viral interactors set: those known to interact with viral proteins that are immunomodulators (proteins known to target the host immune system and modulate its response, obtained from Pichlmair, A. *et al.* ^39^). The distributions for each set were compared using empirical p-values.

### Statistical analysis

Statistical analyses were done with R version 3.3.2. Empirical p-values were calculated in cases where two compared groups had a large difference in their sizes, by randomly subsampling the larger group to the size of the smaller group 10,000 times, and comparing the resulting median (or mean) of each of these subsets to the median (or mean) of the compared smaller group. Throughout the paper, *** denotes p-value<0.001, ** p-value<0.01, * p-value<0.05.

